# Sirt1 overexpression attenuates high-fat diet induced aortic stiffening

**DOI:** 10.1101/2020.10.28.358804

**Authors:** Venkateswara R. Gogulamudi, Daniel R. Machin, Grant Henson, Anthony J. Donato, Lisa A. Lesniewski

## Abstract

Increased arterial stiffness is a cardiovascular disease risk factor in the setting of advancing age and high-fat (HF) diet induced obesity. Increases in large artery stiffness, as measured by pulse wave velocity (PWV), occur within 8 weeks of HF feeding in mice. Sirtuin-1 (Sirt1), a NAD-dependent deacetylase, regulates cellular metabolic activity and activation of this protein has been associated with vasoprotection in aged mice. The aim of the present study was to elucidate the effect of global Sirt1 overexpression (Sirt^tg^) on HF diet-induced arterial stiffening. Sirt1 overexpression did not influence PWV in normal chow (NC) fed mice (Sirt^tg^: 263 ± 6 vs WT: 274 ± 7, p=0.28). However, PWV was higher in wild-type (WT) mice (376 ± 22, p<0.04), but not Sirt^tg^ (304 ± 2 cm/s, p=0.07), after 12 weeks of HF diet. Despite no effect of Sirt1 overexpression on aortic collagen content in NC (p=0.71), aortic elastin content was higher in Sirt^tg^ mice compared with WT mice fed NC diet (P<0.05). Surprisingly, despite increased arterial stiffness, collagen content was lower (p<0.02) and elastin content was unchanged (p=0.05) in the aortas of WT mice after HF. Neither collagen (p=0.18) nor elastin content (p=0.56) were impacted by HF diet in the Sirt^tg^ mice. Likewise, there was no difference in wall thickness in NC (Sirt^tg^: 40.7 ± 2 vs WT: 41.6 ± 2, p= 0.78). However, wall thickness was higher in mice WT mice fed a HF diet (51.7 ± 2, p<0.01) and there was no difference in Sirt^tg^ mice after HF diet (p=0.66). Similarly, there was no difference in wall-to-lumen ratio in mice fed NC diet (Sirt^tg^: 0.08 vs WT: 0.08, p=0.48) was higher in HF diet fed WT mice (p<0.01), though, HF diet was associated with a higher wall-to-lumen ratio in WT (0.11, p<0.01), but not different in Sirt^tg^ mice fed HF diet (0.08, p=0.59). These findings demonstrate a vasoprotective effect of Sirt1 overexpression that limits increases in arterial stiffness and protects against alterations in vessel morphology in response to HF feeding. As such, activation of Sirt1 may be a novel therapeutic target to prevent elevated CVD risk associated with HF-induced aortic stiffening.

## Introduction

Western diet and obesity are associated with cardiovascular disease (CVD), the leading cause of morbidity and mortality worldwide [1]. Arterial dysfunction, characterized by endothelial dysfunction and increased large artery stiffness, occurs with high fat (HF) feeding/obesity [1-3] and is an independent risk factor for CVD [4, 5]. Indeed, long term ingestion of HF diet leads to increases in large artery stiffness resulting in increased pulse wave velocity (PWV) [6, 7] and hypertension [3]. These outcomes are associated with morphological changes in the vasculature such as alterations in vessel diameter and wall thickness [8] and/or changes in structural components of the arterial wall, such as collagen and elastin [7]. With the high prevalence of excessive dietary fat consumption [6] and obesity [9], it is essential to understand the mechanisms underlying increases in arterial stiffness and to develop strategies to combat increased CVD risk in these populations.

Sirtuin-1 (Sirt1) belongs to the NAD^+^-dependent class III histone deacetylases [10]. In mammals, seven sirtuin homologues designated as Sirt1-Sirt7 have been identified, all seven sirtuins share a common catalytic core [11]. Sirtuins have been implicated in regulation of various biological systems in association with caloric restriction, metabolism, aging, cancer, cell differentiation, chromosomal stability, stress resistance, inflammation, DNA repair, tissue fibrosis, mitochondrial biogenesis and apoptosis [12-15]. Sirt1 plays a regulatory role, deacetylation of non-histone proteins such as endothelial nitric oxide (eNOS), Forkhead Box O (FOXO), liver x receptor (LXR), and tumor protein p53 (P53) and peroxisome proliferator-activated receptor-gamma coactivator alpha (PGC1 alpha) [16-20]. Importantly, recent studies have also demonstrated a detrimental effect of reduced Sirt1 on endothelial function that was associated with a reduced activation of eNOS and impaired NO production in aged mice [21]. In contrast, pharmacological activation of Sirt1 has been shown to reverse endothelial dysfunction that was associated with a reduction of both ROS production and inflammation in aged mice [22]. However, the effects of Sirt1 overexpression on arterial stiffness in the setting of HF diet is not known.

In the present study, we tested the hypothesis that Sirt1 overexpression will attenuate HF diet-induced aortic stiffening. To do so, we assessed in-vivo aortic stiffness by PWV in a global Sirt1 overexpressing transgenic mouse model after the consumption of either NC or HF diet for 12 weeks as previously described [7]. In addition, we examined the impact of Sirt1 overexpression on changes in body mass and blood pressure as well as examined the impact of Sirt1 overexpression on HF diet associated alterations in the key arterial structural proteins, collagen and elastin, and vascular morphology.

## Material and Methods

### Animals

Male and female Sirt1 transgenic (TG) and wild type (WT) littermate control mice on C57BL/6 background were generated from breeding colonies at the animal facility of VA Medical Center, Salt Lake City [23]. All the mice were housed at the animal facility of VA Medical Center, Salt Lake City at 23-25°C on a 12 h light/dark cycle and fed water and food ad libitum. 5-6months old male and female mice weighing around 25-30 g were used in this study. Mice were fed a normal (NC) chow (16% kcal from fat, 55% carbohydrate, 29% protein, normal chow TD#8640 Harlan Teklad 22/5 Standard Rodent Chow) or a commercially available high fat (HF) chow (41% kcal from fat (41% saturated/total fat, 41% from carbohydrate, 18% protein, Harlan Adjusted Fat diet #TD.96132) ad libitum for 12 weeks prior to sacrifice [7, 21]. When possible, outcomes were assessed pre (NC) and post HF diet. All animal procedures conformed to the *Guide to the Care and Use of Laboratory Animals: Eighth Edition [24]*, and were approved by the Salt Lake City Veteran’s Affairs Medical Center and University of Utah use Committees.

### Aortic pulse wave velocity (PWV)

Aortic PWV was measured before and 12 weeks after initiation of HF as described previously [7, 25]. Briefly, mice were anesthetized with 2% isoflurane in a closed compartment anesthesia machine (V3000PK, Parkland Scientific, Coral Springs, FL) for ~1-3 min. Throughout the procedure anesthesia was maintained via a nose cone and mice were secured in a supine position on a heated board (~35°C) to maintain body temperature. PWV was measured with 20-MHz Doppler probes (Indus Instruments, Webster, TX) at the transverse aortic arch and ~4 cm distal at the abdominal aorta and collected using WinDAQ Pro+ software (DataQ Instruments, Akron, OH). Absolute pulse arrival times were indicated by the sharp upstroke, or foot, of each velocity waveform analyzed with WinDAQ Waveform Browser (DataQ Instruments, Akron, OH) and distance between the probes was measured using a scientific caliper. Velocities were then calculated as the quotient of the separation distance (~3 cm) and difference in absolute arrival times.

### Blood Pressure

Blood pressure was measured in NC and HF fed WT and Sirt^tg^ mice via non-invasive tail-cuff (CODA System, Kent Scientific, CT) technique as described previously [25]. Briefly, mice were acclimated to the restrainers and then placed on a heating unit to reach a steady body (skin and tail) temperature (37°C) in a quiet room. Systolic blood pressure (SBP) was calculated from the average of 15 recordings.

### Vessel morphology

After HF feeding mice were euthanized and aortas were collected and prepared for histological analyses. All mice were euthanized by exsanguination via cardiac puncture under isoflurane anesthesia. Thoracic aortas were excised and placed in physiological saline at 4°C. With the aid of a dissecting microscope, perivascular tissues were cleared and ~2 millimeter (mM) rings were cut from the descending thoracic aorta distal to the greater curvature. The rings were embedded in Optimal Cutting Temperature (OCT) medium. From each mouse, three frozen sections of aorta (8 micron) were made using a cryostat (Thermo Scientific) and mounted on a glass slide for collagen and elastin quantification. Collagen staining was performed using picrosirius red, wall thickness was calculated as the radius of the outer border of the medial layer minus radius of the lumen area and wall to lumen ratio was calculated as wall thickness divided by lumen diameter quantified by ImageJ (NIH, Bethesda, MA) as described previously [25, 26]. Elastin was quantified by Verhoff’s Van Geison stain as described previously [27]. An 8-bit grayscale was used for densitometric quantification of elastin content with ImageJ.

### Western blotting

Thoracic aortas from NC and HF fed WT and Sirt^tg^ mice were dissected, cleared of the perivascular fat and stored in liquid nitrogen. Aortic lysates were prepared in RIPA buffer by physical disruption with a homogenizer and supernatants were collected. After the total protein quantification, equal amounts of protein (20 ug) were loaded in 4-12% - Criterion XT Bis-Tris protein gels (Bio-Rad, CA) and transferred to nitrocellulose membranes. The membranes were blocked with 1% BSA with TBST before incubation with Sirt1 primary antibody (1:200, Santa Cruz, CA) at 4°C overnight. Membranes were then washed 3 times with TBST and incubated with secondary antibody for 1hr at room temperature. Blots were developed by super signal ECL reagent and bands were visualized using a gel documentation system. To normalize the protein loading differences, vinculin protein was used as housekeeping protein expression and data was normalized to the mean of the WT group.

### Statistics

To determine if differences existed between genotypes and diets for terminal measures, a two-way ANOVA was conducted. To determine if differences existed between genotypes and diets for in vivo measures, a two-way repeated measures ANOVA was performed. All the data are represented as mean ± SEM, significance was set as p<0.05.

## Results

### Animal characteristics

Sirt1 protein expression in the thoracic aorta was higher in NC fed Sirt^tg^ compared with WT mice (p<0.01, **Fig.1A**). There was no effect of Sirt1 overexpression on body, heart, liver, spleen soleus, gastrocnemius, quadriceps or WAT mass in NC fed mice. WT mice had higher body mass (p<0.05) and lower heart, liver, soleus, and quadriceps mass (all p<0.05) after HF diet compared with NC fed WT mice. Similarly, compared with NC fed mice, the mass of the heart and quadriceps muscle was lower (p<0.05) but gastrocnemius muscle mass was higher (p<0.05, Table 1) in HF fed Sirt1^tg^ mice.

**Table 1.** Body, heart, liver, spleen, soleus, gastrocnemius, quadriceps muscle, and epididymal white adipose tissue (WAT) mass in WT and Sirt^tg^ normal chow (NC) and high-fat (HF) -fed mice. Values are mean ± SEM \**P* < 0.05 versus WT-NC. \**P* < 0.05 versus TG-NC.

|  | WT - NC | WT - HF | Sirt<sup>tg</sup> - NC | Sirt<sup>tg</sup> - HF |
| --- | --- | --- | --- | --- |
| **Male : female N** | 10:4 | 6:7 | 11:6 | 8:8 |
| **Age** | 6.0 ± 0.3 | 8.0 ± 0.5 | 6.0 ± 0.2 | 7.7 ± 0.2<sup>#</sup> |
| **Body mass (g)** | 27.3 ± 1 | 33.6 ± 1* | 27.3 ± 1 | 27.6 ± 1 |
| **Heart mass (g)** | 0.168 ± 0 | 0.121 ± 0* | 0.160 ± 0 | 0.125 ± 0<sup>#</sup> |
| **Liver mass (g)** | 1.916 ± 0 | 1.509 ± 0* | 1.737 ± 0 | 1.483n ± 0 |
| **Spleen mass (g)** | 0.082 ± 0 | 0.086 ± 0 | 0.098 ± 0 | 0.076 ± 0 |
| **Soleus mass (g)** | 0.018 ± 0 | 0.011 ± 0* | 0.014 ± 0 | 0.010 ± 0 |
| **Gastrocnemius mass (g)** | 0.243 ± 0 | 0.152 ± 0* | 0.147 ± 0 | 0.210 ± 0<sup>#</sup> |
| **Quadriceps mass (g)** | 0.267 ± 0 | 0.185 ± 0* | 0.226 ± 0 | 0.182 ± 0<sup>#</sup> |
| **WAT mass (g)** | 0.800 ± 0 | 0.457 ± 0 | 0.486 ± 0 | 0.562 ± 0 |

**Figure. 1.**
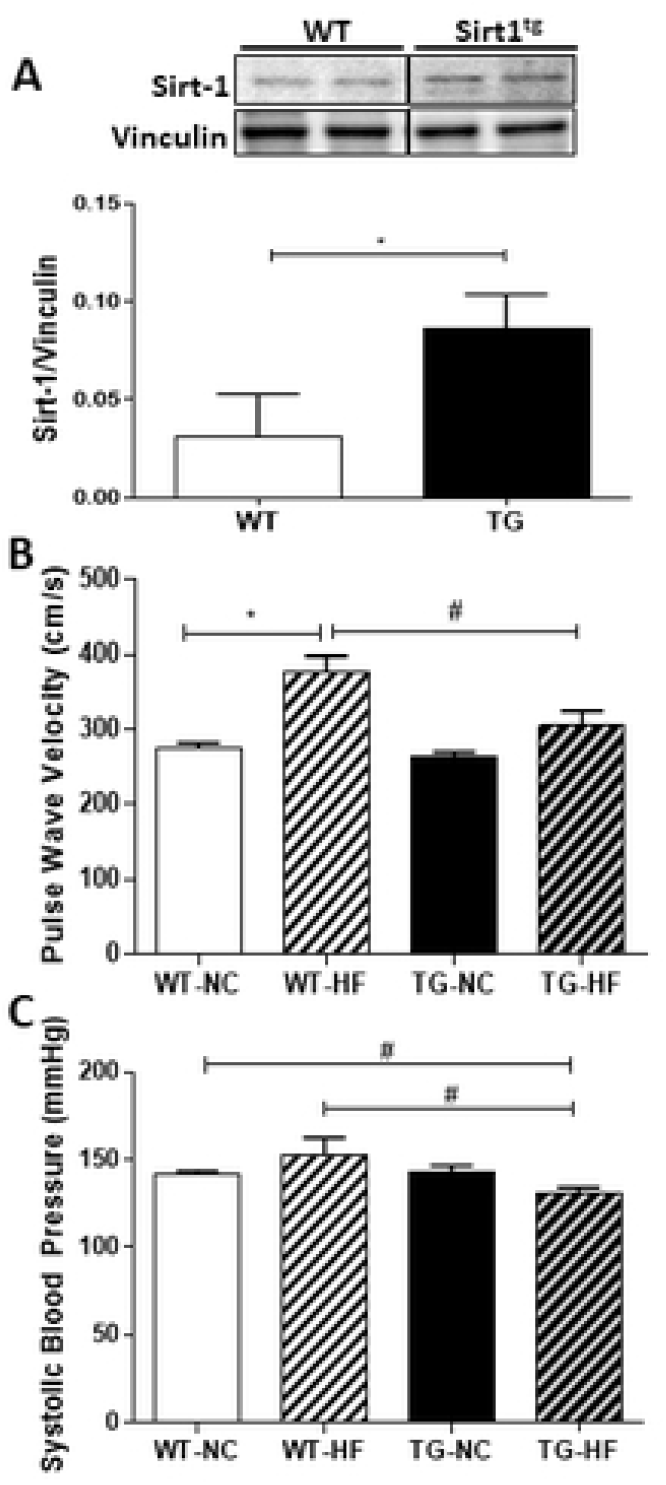
(**A**) Sirt1 protein expression measured by western blotting, in thoracic aorta excised from WT and Sirt1^tg^ mice (**B**) Aortic pulse wave velocity (PWV) assessed in WT and Sirt^tg^ littermates before (PRE) and following 12 weeks of high fat diet (POST) (**C**) Systolic blood pressure (SBP)assessed in WT and Sirt^tg^ littermates before (PRE) and following 12 weeks of high fat diet (POST) *denotes a significant PRE/POST difference within genotype,^#^ denotes a significant difference WT Vs Sirt^tg^ genotype. Data are mean ± SEM.

### Aortic Stiffness

Sirt1 overexpression did not influence aortic stiffness, assessed by aortic PWV, in NC fed WT mice (Sirt^tg^: 263 ± 6 vs WT:, p=0.28). However, the effect of HF diet to increase aortic stiffness in WT mice (274 ± 7 vs. 376 ± 22, p<0.04) was lost in the Sirt^tg^ mice (263 ± 6 vs, 304 ± 2 cm/s, p=0.07) (**Fig.1B**). Anesthetized heart rate was similar between groups (WT-NC: 341± 30; WT-HF: 397±30; Sirt^tg^-NC: 364±31 Sirt’g-HF: 381±27 beats/min, p>0.05).

### Blood pressure

Sirt1 overexpression did not impact systolic blood pressure (SBP) in NC fed mice (Sirt^tg^: 142 ± 2 vs WT: 141 ± 3 mmHg, p=0.82). Although there were no significant effects of HF diet on SBP in either the WT or Sirt^tg^ (130 ± 3, p=0.14 Vs NC), SBP tended to increase in WT (152 ± 9, P=0.14) after HF diet, resulting in a significantly lower SBP in the HF fed Sirt^tg^ compared to diet matched WT mice (p<0.05) (**Fig.1C**) after HF diet.

### Collagen and elastin content

There was no effect of Sirt1 overexpression on aortic collagen content in NC fed mice (p=0.71). Despite increased aortic stiffness after HF diet, collagen content was lower in the aortas of WT mice compared to NC fed mice (p<0.006). There was no difference in collagen content in response to HF feeding in the Sirt^tg^ mice (p=0.07). Surprisingly, we found that collagen content was higher in the aortas of HF fed Sirt^tg^ mice compared to HF fed WT mice (p<0.02) despite lower PWV (**Fig. 2A**), suggesting that alterations in collagen are not the primary determinant of in vivo arterial stiffness in this model. Despite no difference in PWV between groups, elastin content was higher in Sirt^tg^ compared with WT mice fed NC diet (p<0.05). There was no difference in elastin content after HF diet in either the WT or Sirt^tg^ mice (**Fig. 2B**), again suggesting that changes in these primary structural components is not the primary determinant of in vivo changes in aortic stiffness.

**Figure 2.**
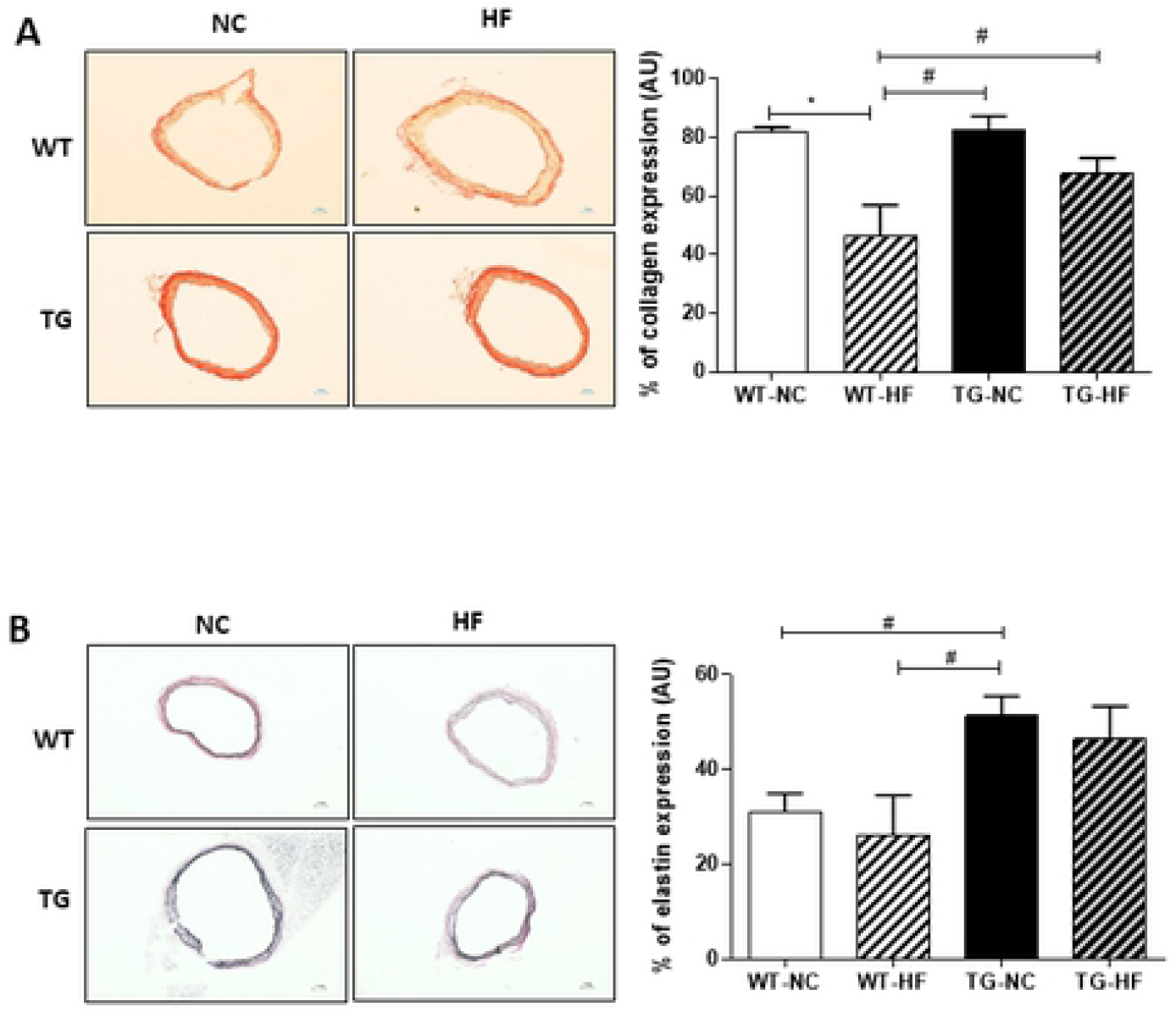
**(A)**. Total collagen content, measured by picrosirius red stain, in thoracic aorta excised from WT and Sirt^tg^ mice fed either normal chow (NC) or high fat diet (HF) for 12 weeks. **(B)**Total elastin content, measured by Verheoff’s Van Geisson stain, in thoracic aorta excised from WT and Sirt^tg^ mice fed either normal chow (NC) or high fat diet (HF) for 12 weeks. *denotes a significant PRE/POST difference within genotype, ^#^ denotes a significant difference WT Vs Sirt^tg^ genotype. Data are mean ± SEM.

### Aortic vessel characteristics

There was no difference in wall thickness between NC fed Sirt^tg^ and WT mice (p>0.78). However, there was higher wall thickness after HF diet in WT mice (p<0.01), but not impacted the Sirt^tg^ mice (p=0.66). There was no significant effect of NC diet on wall to lumen ratio in either the WT or Sirt^tg^ mice (p>0.48). Although there was a significantly higher wall to lumen ratio after HF diet in WT mice (p<0.02), there was no difference in Sirt^tg^ mice after HF diet (p=0.66). (**Fig. 3**).

**Figure 3.**
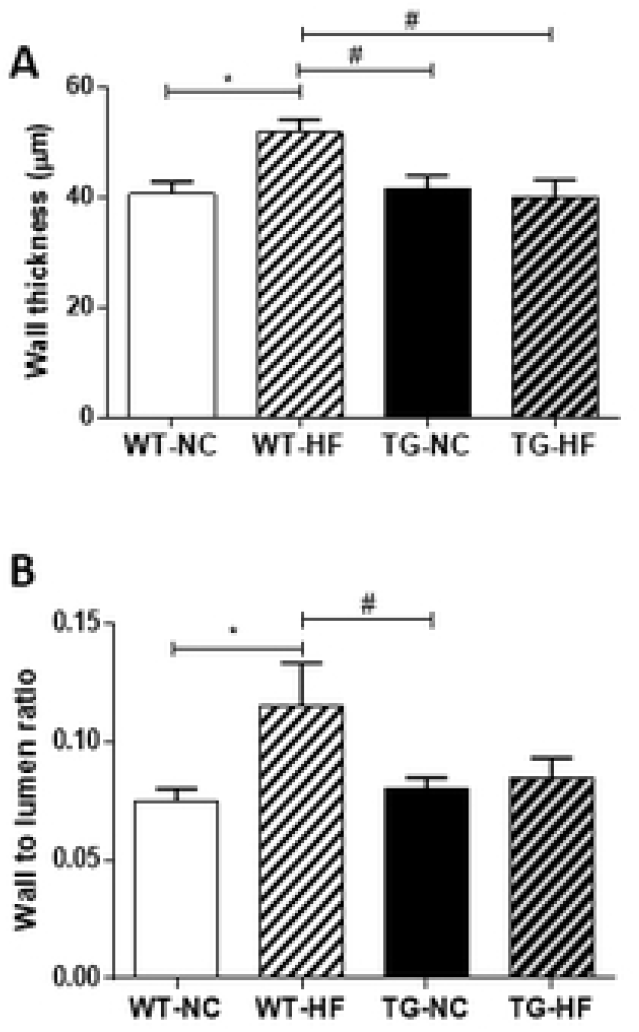
Wall thickness (**A**), wall to lumen ratio (**B**) in thoracic aorta from WT and Sirt^tg^ mice fed either normal chow (NC) or high fat diet (HF) for 12 weeks. *denotes a significant PRE/POST difference within genotype, ^#^ denotes a significant difference WT Vs Sirt^tg^ genotype. Data are mean ± SEM

## Discussion

The novel findings of the present study are that Sirt1 overexpression protects against HF diet–induced large artery stiffening and that this is associated with a protection against arterial wall remodeling. Specifically, we found that Sirt1 overexpression lowers SBP, and prevents an increase in wall thickness, wall-to-lumen ratio in response to HF feeding, but this protection appears to be independent of changes in the structural components of the arterial wall, such as collagen and elastin. Our findings suggest that Sirt1 activation may be a viable therapeutic strategy to combat arterial stiffening in the face of obesity/high fat diet consumption.

Previous studies have demonstrated that Sirt1 can play a protective role against experimental hypertension. For example, vascular remodeling in response to angiotensin II induced hypertension was attenuated in a genetic model of vascular smooth muscle Sirt1 overexpression [27]. Likewise, smooth muscle cell specific Sirt1 overexpression and pharmacological activation of Sirt1 in a smooth muscle specific Sirt1 knockout model have been shown to protect against HF and high sucrose diet induced increases in arterial stiffness measured by PWV [28] and this vasoprotection was associated with reductions in vascular inflammation [28, 29]. In the present study, we used a global Sirt1 overexpression mouse model and examined aortic stiffness and structure to provide a more complete understanding of the impact of Sirt1 on HF associated increases in large artery stiffening. In agreement with the known benefits of Sirt1 overexpression, we observed that HF induced aortic stiffness was attenuated in Sirt1 transgenic mice compared with WT mice. As increases in aortic stiffness are an independent risk factor for CVD, our data suggests that interventions to increase Sirt1 activation may be efficacious to reduce arterial stiffening in obesity. Similar to aortic stiffness, Sirt1 overexpression also prevented the increase in SBP observed in after HF diet. This is an agreement with the previous studies, in which vascular smooth muscle specific (VSMC) and whole body Sirt1 overexpression protected against long term (8 months) diet-induced aortic stiffening and further argues for the vasoprotective effects of Sirt1 [29].

The results of the present study expand upon this previous report by demonstrating that global overexpression of Sirt1 protects against short term (12 weeks) HF diet-induced increases in large artery stiffness and extends the previous study by demonstrating that this protective effect is associated with alterations in vessel morphology. Although we found differential changes in collagen and elastin in response to either Sirt1 overexpression and HF feeding, the direction of these changes is such that they are not likely to contribute to the observed functional changes in arterial stiffness. We, and others, have demonstrated that arterial stiffening ensues within 8-12 weeks of HF feeding in mice and that this is associated with vascular remodeling [7, 29] and structural modifications in the arterial wall [25, 30]. Here, we find that Sirt1 overexpression protects against HF diet-induced medial hypertrophy, ameliorating the increase in the wall thickness and wall-to-lumen ratio observed in response to HF diet in WT mice. The extent to which these effects are facilitated by antioxidant and anti-inflammatory activity as suggested by the smooth muscle specific pharmacological activation of Sirt1 [29] is unclear but requires further elucidation.

Here, we find that the Sirt1 overexpression did not impact collagen content as expected from the changes in PWV observed. Indeed, we found no difference in collagen content in NC fed mice, and a greater collagen content in the aortas of HF fed Sirt^tg^ mice compared to WT mice, suggesting that collagen content is not a primary determinant of the changes in the arterial stiffness measured in vivo by PWV. Furthermore, we find that Sirt1 overexpression-mediated prevention of arterial stiffening in response to HF diet is associated with a maintenance of higher elastin content compared to WT mice regardless of diet. This is largely consistent with previous findings that Sirt1 activation by nicotinamide mononucleotide supplementation lowers aortic stiffness and maintains the elastin and collagen content [29]. Thus, although maintenance of higher elastin content in the Sirt1 transgenic mice even in the face of HF diet is one potential contributor to the amelioration of HF diet induced arterial stiffening, it is not likely the only factor as HF diet did not significantly reduce elastin content in the WT mice. While it is unclear how the present study’s findings of increased arterial stiffness and lower collagen content in HF fed WT mice can coincide, changes in in vivo vascular tone is one possibility. While alterations in collagen and elastin may impact the mechanical properties of the vascular wall, impacting the material stiffness of the artery; but these changes may be masked in vivo by changes in arterial tone. Indeed, arterial stiffness is strongly affected by vascular tone, which is itself modified by endothelial function. In vivo, endothelial cell function can be modified by mechanostimulation, cell stretch, changes in calcium signaling and by paracrine mediators including angiotensin II, oxidant stress, endothelin and nitric oxide [31]. Although the role of altered vascular tone in the modulation of arterial stiffening by Sirt1 overexpression should be further explored, we and others have demonstrated that endothelial Sirt1 has a regulatory effect on vasodilation and vascular tone [16, 21, 32] that may explain the decreases in HF diet induced arterial stiffening in the transgenic mice.

In the setting of HF diet-induced obesity, elevated blood pressure has also been implicated in increased arterial stiffness via increases in pulsatile pressure and pulsatile aortic wall stress that can accelerate elastin degradation [33, 34]. Here, it is unclear if an attenuation of the hypertensive response to HF feeding is an underlying mechanism for the protection against increases in aortic stiffening. Although SBP was lower in Sirt1 transgenic mice compared to WT mice after HF diet which corresponded to a lower aortic pulse wave velocity, this was not accompanied by a significant increase in SBP in WT mice in response to HF feeding. Taken together our data indicates that whole body Sirt1 overexpression is involved in the prevention of HF induced vascular stiffening that is associated with prevention of morphological and structural changes.

Mice fed a HF diet mimic human vascular and metabolic diseases and allow for the investigation of mechanisms underlying vascular consequences of obesity. Previously, we demonstrated arterial stiffness increases in the B6D2F1 mouse model in response to chronic HF feeding and aging [7]. Although increased arterial stiffness is an independent risk factor for CVD, there is no specific treatment available to humans. Activation of Sirt1 may be a potential therapeutic target to treat elevations in arterial stiffness with either obesity or aging, as Sirt1 is abundantly expressed in vasculature and is involved in the maintenance of vascular homeostasis by providing protection against detrimental vascular tissue remodeling, atherosclerosis and endothelial senescence [32, 35, 36], as well as in the regulation of vasodilation and vascular tone [10, 19, 35]. Interestingly, Sirt1 is modulated in states of increased CVD risk such as aging and obesity. Indeed, Sirt1 expression decreases in endothelium with aging [21, 37] and HF diet induces inflammation triggered cleavage of Sirt1 in adipose tissue which promotes metabolic dysfunction [38]. In this present study we demonstrated that overexpressing Sirt1 protects mice against arterial stiffening in response to HF diet. Our study suggests that Sirt1 activation may be a novel therapeutic target for treating obesity-associated increases in arterial stiffness and deleterious arterial remodeling.

## Summary

The novel findings of this study reveals that Sirt1 overexpression prevents the aortic stiffening in response to HF feeding and suggest that this is associated with an increased elastin content of the aorta. We further demonstrate that Sirt1 overexpression may contribute to a lowering of SBP in the face of HF diet. The findings suggest that Sirt1 plays an important protective role against HF diet induced large artery stiffening and may be a therapeutic target in the management of arterial stiffening and blood pressure in a metabolic disease population.

## Acknowledgements

All experiments were performed in the Translational Vascular Physiology Laboratory at the University of Utah and Veteran’s Affairs Medical Center-Salt Lake City Geriatrics Research Education and Clinical Center. The authors have no conflicts of interest to disclose.

This work was supported by National Institutes of Health awards: R01 AG060395, R01 AG050238, R01 AG048366, K02 AG045339, R00 AT010017 and Veteran’s Affairs Merit Review Awards I01 BX002151 and I01 BX004492 from the United States (U.S.) Department of Veterans Affairs Biomedical Laboratory Research and Development Service. The contents do not represent the views of the U.S. Department of Veterans Affairs, the National Institutes of Health or the United States Government.

## References

1. Benjamin, E.J., et al., Heart Disease and Stroke Statistics-2019 Update: A Report From the American Heart Association. Circulation, 2019. 139(10): p. e56–e528.

2. Donato, A.J., et al., TNF-alpha impairs endothelial function in adipose tissue resistance arteries of mice with diet-induced obesity. Am J Physiol Heart Circ Physiol, 2012. 303(6): p. H672–9.

3. Weisbrod, R.M., et al., Arterial stiffening precedes systolic hypertension in diet-induced obesity. Hypertension, 2013. 62(6): p. 1105–10.

4. Scuteri, A., et al., Arterial stiffness is an independent risk factor for cognitive impairment in the elderly: a pilot study. J Hypertens, 2005. 23(6): p. 1211–6.

5. Lacy, P.S., et al., Increased pulse wave velocity is not associated with elevated augmentation index in patients with diabetes. J Hypertens, 2004. 22(10): p. 1937–44.

6. Santana, A.B., et al., Effect of high-fat diet upon inflammatory markers and aortic stiffening in mice. Biomed Res Int, 2014. 2014: p. 914102.

7. Henson, G.D., et al., Dichotomous mechanisms of aortic stiffening in high-fat diet fed young and old B6D2F1 mice. Physiol Rep, 2014. 2(3): p. e00268.

8. Dantas, A.P., et al., Western diet consumption promotes vascular remodeling in non-senescent mice consistent with accelerated senescence, but does not modify vascular morphology in senescent ones. Exp Gerontol, 2014. 55: p. 1–11.

9. Wildman, R.P., et al., Measures of obesity are associated with vascular stiffness in young and older adults. Hypertension, 2003. 42(4): p. 468–73.

10. Tajbakhsh, N. and E.M. Sokoya, Regulation of cerebral vascular function by sirtuin 1. Microcirculation, 2012. 19(4): p. 336–42.

11. Lu, G., et al., Role and Possible Mechanisms of Sirt1 in Depression. Oxid Med Cell Longev, 2018. 2018: p. 8596903.

12. Michan, S. and D. Sinclair, Sirtuins in mammals: insights into their biological function. Biochem J, 2007. 404(1): p. 1–13.

13. Ozawa, Y., et al., Retinal aging and sirtuins. Ophthalmic Res, 2010. 44(3): p. 199–203.

14. Brochier, C., et al., Specific acetylation of p53 by HDAC inhibition prevents DNA damage-induced apoptosis in neurons. J Neurosci, 2013. 33(20): p. 8621–32.

15. Wojcik, M., K. Mac-Marcjanek, and L.A. Wozniak, Physiological and pathophysiological functions of SIRT1. Mini Rev Med Chem, 2009. 9(3): p. 386–94.

16. Mattagajasingh, I., et al., SIRT1 promotes endothelium-dependent vascular relaxation by activating endothelial nitric oxide synthase. Proc Natl Acad Sci U S A, 2007. 104(37): p. 14855–60.

17. Brunet, A., et al., Stress-dependent regulation of FOXO transcription factors by the SIRT1 deacetylase. Science, 2004. 303(5666): p. 2011–5.

18. Li, X., et al., SIRT1 deacetylates and positively regulates the nuclear receptor LXR. Mol Cell, 2007. 28(1): p. 91–106.

19. Vaziri, H., et al., hSIR2(SIRT1) functions as an NAD-dependent p53 deacetylase. Cell, 2001. 107(2): p. 149–59.

20. Rodgers, J.T., et al., Nutrient control of glucose homeostasis through a complex of PGC-1alpha and SIRT1. Nature, 2005. 434(7029): p. 113–8.

21. Donato, A.J., et al., SIRT-1 and vascular endothelial dysfunction with ageing in mice and humans. J Physiol, 2011. 589(Pt 18): p. 4545–54.

22. Gano, L.B., et al., The SIRT1 activator SRT1720 reverses vascular endothelial dysfunction, excessive superoxide production, and inflammation with aging in mice. Am J Physiol Heart Circ Physiol, 2014. 307(12): p. H1754–63.

23. Banks, A.S., et al., SirT1 gain of function increases energy efficiency and prevents diabetes in mice. Cell Metab, 2008. 8(4): p. 333–41.

24. in Guide for the Care and Use of Laboratory Animals, th, Editor. 2011: Washington (DC).

25. Donato, A.J., et al., Life-long caloric restriction reduces oxidative stress and preserves nitric oxide bioavailability and function in arteries of old mice. Aging Cell, 2013. 12(5): p. 772–83.

26. Fleenor, B.S., et al., Arterial stiffening with ageing is associated with transforming growth factor-beta1-related changes in adventitial collagen: reversal by aerobic exercise. J Physiol, 2010. 588(Pt 20): p. 3971–82.

27. Raub, C.B., et al., Linking optics and mechanics in an in vivo model of airway fibrosis and epithelial injury. J Biomed Opt, 2010. 15(1): p. 015004.

28. Gao, P., et al., Overexpression of SIRT1 in vascular smooth muscle cells attenuates angiotensin II-induced vascular remodeling and hypertension in mice. J Mol Med (Berl), 2014. 92(4): p. 347–57.

29. Fry, J.L., et al., Vascular Smooth Muscle Sirtuin-1 Protects Against Diet-Induced Aortic Stiffness. Hypertension, 2016. 68(3): p. 775–84.

30. Machin, D.R., et al., Lifelong SIRT-1 overexpression attenuates large artery stiffening with advancing age. Aging (Albany NY), 2020. 12(12): p. 11314–11324.

31. Zieman, S.J., V. Melenovsky, and D.A. Kass, Mechanisms, pathophysiology, and therapy of arterial stiffness. Arterioscler Thromb Vasc Biol, 2005. 25(5): p. 932–43.

32. Wang, Y., Y. Liang, and P.M. Vanhoutte, SIRT1 and AMPK in regulating mammalian senescence: a critical review and a working model. FEBS Lett, 2011. 585(7): p. 986–94.

33. Mitchell, G.F., Arterial stiffness and hypertension: chicken or egg? Hypertension, 2014. 64(2): p. 210–4.

34. Safar, M.E., S. Czernichow, and J. Blacher, Obesity, arterial stiffness, and cardiovascular risk. J Am Soc Nephrol, 2006. 17(4 Suppl 2): p. S109–11.

35. Potente, M. and S. Dimmeler, Emerging roles of SIRT1 in vascular endothelial homeostasis. Cell Cycle, 2008. 7(14): p. 2117–22.

36. Bai, B., et al., Endothelial SIRT1 prevents adverse arterial remodeling by facilitating HERC2-mediated degradation of acetylated LKB1. Oncotarget, 2016. 7(26): p. 39065–39081.

37. Zu, Y., et al., SIRT1 promotes proliferation and prevents senescence through targeting LKB1 in primary porcine aortic endothelial cells. Circ Res, 2010. 106(8): p. 1384–93.

38. Chalkiadaki, A. and L. Guarente, High-fat diet triggers inflammation-induced cleavage of SIRT1 in adipose tissue to promote metabolic dysfunction. Cell Metab, 2012. 16(2): p. 180–8.

